# Potential Role of Nociceptin/Orphanin FQ in the Progression of Multiple Sclerosis

**DOI:** 10.64898/2026.07.02.736158

**Authors:** Joseph C. Baker, Caitlin E. Paisley, Molly A. Poore, John W. Bigbee, Unsong Oh, Carmen Sato-Bigbee

## Abstract

We showed before that the endogenous peptide Nociceptin blocks the premature differentiation of oligodendrocytes (OLGs), preventing untimely precocious myelination in the developing brain. Consistent with this early function, Nociceptin brain expression is developmentally regulated, sharply decreasing with the initiation and progression of myelination. However, we now found that at difference with controls and relapsing-remitting multiple sclerosis (RRMS), Nociceptin levels are highly elevated in cerebrospinal fluid from patients with the most severe progressive MS (PMS) forms. This questioned whether Nociceptin early developmental effects could be latter recapitulated, interfering with remyelination in PMS. This possibility was tested by inducing experimental autoimmune encephalomyelitis in older mice, at an age equivalent to that with increased risk of RRMS transition into PMS. Older animals develop persistently highly debilitating clinical symptoms, and display both brain and spinal cord demyelination. Importantly, these mice exhibit elevated brain Nociceptin levels, and their treatment with an antagonist of the Nociceptin receptor (NOR) elicits a regression of clinical scoring that is accompanied by higher ratios of OLGs/OLG progenitor cells, increased myelination, and reduction of reactive astrocytes. These findings suggest that Nociceptin may be a crucial player in the age-related progression of MS; interfering with OLG maturation and remyelination, and perhaps further exacerbating neurological dysfunction by targeting astrocyte populations. The upregulation of Nociceptin secretion by human astrocytes in response to proinflammatory cytokines, also points to this peptide as a mediator of microglia-astrocyte interactions supporting MS progression with aging. NOR may offer a novel pharmacological target for ameliorating the devastating effects of MS progression.

## INTRODUCTION

Myelin, the complex multilamellar structure wrapping around axons, maximizes neuronal communication by working as an insulator that facilitates the rapid “saltatory” propagation of electrical impulses ^1^. Furthermore, myelin and the oligodendrocytes (OLGs) that make this specialized membrane in the central nervous system (CNS) are also critical regulators of plasticity; playing important roles in the protection of neurons and regulation of axonal growth; participating in glial-neuronal interactions; and integrating white matter networks that are central to neurodevelopment, behavior and cognition ^2–7^. Both myelin and OLGs are damaged in multiple sclerosis (MS), an inflammatory demyelinating disease that is the leading cause of neurological disability affecting individuals in their most productive years of life ^8^. More than two million people worldwide and nearly 1 million in the United States alone, are estimated to be affected by MS ^9–11^. MS patients exhibit variable symptoms including visual problems, limb weakness, sensory loss, brainstem dysfunction, ataxia, incontinence, sexual dysfunction, fatigue, and cognitive decline ^12; 13^. Underlying these problems is the presence of neurodegeneration caused by local inflammation, multifocal demyelination, astrogliosis, neuro-axonal degeneration, and blood brain barrier dysfunction ^14^. Most MS patients are diagnosed between the ages of 20 and 40, with the majority of cases exhibiting an initial clinical course representative of relapsing remitting MS (RRMS), in which periods of demyelination and worsening neurological symptoms lasting from days to weeks alternate with phases of remyelination and symptoms relief. Only a small number of patients suffers of primary-progressive MS (PPMS) characterized by continual disability increase from disease onset ^15^. Importantly, aging plays a very crucial role in not only the worsening of PPMS, but also in the negative evolution of about 50-80% of the initially milder RRMS cases into more severe secondary-progressive MS (SPMS) ^16–20^. Thus, most current treatments for MS are focused on ameliorating the symptoms of RRMS and prevent or delay the evolution of disease into SPMS. Unfortunately, MS progression remains elusive and there is a lack of effective treatments for SPMS and PPMS (both forms referred from now on in this paper as PMS). Underlying the severity of these PMS forms is the presence of continuing myelin loss and failure of remyelination by insufficient generation of mature OLGs. These cells originate from immature OLG progenitor cells (OPCs) that undergo several rounds of proliferation and steps of differentiation before becoming fully mature myelin-forming OLGs ^21^. This critical sequence of maturational events is also required for remyelination but appears to be disrupted in aging and P-MS. This possibility is supported by different studies that demonstrated that chronically demyelinated lesions display relatively minor decreases of OPC pools but exhibit significantly lower numbers of mature OLGs ^22–25^. This situation is reproduced in animal models of demyelination, showing that while OPCs divide and migrate to regions of myelin loss, their capacity to generate mature OLGs is also reduced by aging ^26^. Therefore, the identification of molecular targets that could be manipulated to stimulate OPC differentiation into mature OLGs capable of myelin regeneration is crucial to the development of therapies to prevent the transition of RRMS into PMS. The results presented here suggest that one of those targets may be Nociceptin/Orphanin FQ (Nociceptin), an endogenous 17-amino acid peptide that specifically binds to the Nociceptin/Orphanin FQ receptor (NOR) and is generated by cleavage of its precursor protein prepronociceptin ^27^. NOR, also known as OPRL-1 or kappa-type 3 opioid receptor, is a G-protein coupled receptor previously implicated in a variety of functions including pain modulation, learning and memory, stress, drug addiction, and the control of glutamate transporter expression in early astrocytes ^28–34^. An inhibitory effect on OLG maturation and myelination triggered by NOR activation was initially suggested by our earlier studies on perinatal exposure to buprenorphine ^35; 36^ , a synthetic opioid used for addiction treatment during pregnancy which at high doses behaves as an NOR agonist. We later found that high Nociceptin levels block the premature differentiation of pre-oligodendrocytes (pre-OLGs) into mature OLGs, preventing untimely precocious myelination that could interfere with axonal elongation and connectivity ^37; 38^. Consistent with this function, Nociceptin levels are developmentally regulated, sharply decreasing with the initiation and progression of myelination ^37^, an expression pattern suggesting that later pathological increases may contribute to impairments in myelin maintenance in the mature brain. We now show that such situation may occur in PMS as Nociceptin levels are highly elevated in the cerebrospinal fluid (CSF) from patients with progressive forms of the disease. The findings presented here using an animal model of MS and aging, suggest that increased local microglial activation and astrocyte reactivity in both PMS and the aging nervous system synergize resulting in elevated Nociceptin concentrations, preventing the maturation of OLGs capable of myelin regeneration. Furthermore, the unexpected re-expression of NOR in reactive astrocytes suggests that MS progression also reflects still unknown Nociceptin effects in different glial populations, perhaps further exacerbating neurological dysfunction.

## MATERIALS AND METHODS

### Materials

Horseradish peroxidase (HRP)-conjugated secondary antibodies including goat anti-rabbit IgG (Cat. No. 111-035-003), rabbit anti-Mouse IgG (Cat. No. 315-035-045), and human-absorbed donkey anti-rabbit IgG (Cat. No. 711-035-152); and all fluorescently labeled secondary antibodies, including Alexa Fluor 594 goat anti-mouse IgG (Cat. No. 115-587-003), Alexa Fluor 594 goat anti-chicken IgG (Cat. No. 103-587-008), Alexa Fluor 488 goat anti-rabbit IgG (Cat. No. 111-547-003), and Alexa Fluor 594 goat anti-rabbit IgG (Cat No. 711-587-003), were from Jackson ImmunoResearch Laboratories (West Grove, PA). Anti-Nociceptin rabbit polyclonal (Cat. PA3-204), anti-Glial fibrillary acidic protein (GFAP) rabbit polyclonal (Cat. No. PA1-10004), anti-NG2 proteoglycan (NG2) rat monoclonal (MA5-24247), and anti-NOR rabbit polyclonal (12970-1-AP) antibodies were from Invitrogen (ThermoFisher, Waltham, MA). The anti-PDGF-R alpha rat polyclonal antibody (Cat. No. 558774) was from BD Biosciences (Milpitas, CA). The anti-Myelin basic protein (MBP) rat monoclonal antibody (Cat. No. MAB386) was from Sigma-Aldrich (St. Louis, MO). The Cy5-conjugated-anti-Myelin regulatory factor (MYRF) rabbit polyclonal antibody (Cat. No. bs-11191R-CY5) was from BIOSS Inc. (Woburn, MA). Invitrogen ProLong Glass Antifade with NucBlue slide mounting solution (Cat. No. P36985), Halt Protease and Phosphatase Inhibitor Cocktail (Cat. No. PI78442), and Super Signal West Dura Extended Duration Substrate for chemiluminescence (Cat. No. 34075) were provided by ThermoFisher Scientific. Myelin oligodendrocyte glycoprotein amino acids 35-55 peptide (MOG_35-55_) (Cat. No. AS-60130) was from ANASPEC (Fremont, CA). Pertussis Toxin (Cat. No. 70323-44-3) was acquired from List Biological Laboratories (Campbell, CA). Mycobacterium Tuberculosis H37Ra (M. Tuberculosis) (Cat. No. 231141) and incomplete Freund’s adjuvant (Cat. No. 263910) were from BD Difco (Detroit, MI). The NOR antagonist BAN-ORL-24[(2R)-1-(phenylmethyl)-N-[3-(spiro[isobenzofuran-1(3H),4’-piperdin]-1-yl)propyl-2-pyrolidinecarboximide] was acquired from APExBIO (Cat. No. A3219) (Houston, TX) and Cayman Chemical (Cat. No.29485) (Ann Arbor, MI). TNF-α (Cat. No. 300-01A-50UG), IL-1β (Cat. No. 200-01B-10UG), and IL-6 (Cat. No. 200-06-20UG) were from PeproTech and provided by ThermoFisher. Tissue Plus Optimal cutting temperature compound (OCT) (Cat. No. 4585) was from Fisher Healthcare (Houston, TX). Nitrocellulose membrane for dot blot and western blot analyses and all electrophoresis reagents were from Bio-Rad (Hercules, CA).

### Animals

C57BL/6J male and female mice were obtained from Jackson Laboratories (Bar Harbor, ME). All animal use was conducted in accordance with the National Institutes of Health (NIH) recommendations and approved by the Virginia Commonwealth University Animal Care and Use Committee (IACUC).

### Human Samples

Cerebrospinal fluid (CSF) samples from de-identified controls and MS patients were obtained from both the Netherlands Brain Bank (Netherlands), and Virginia Commonwealth University (VCU) Brain Tissue Bank/Data Registry housed in the Department of Neurology (VCU School of Medicine).

### Evaluation of Nociceptin secretion by Human Astrocytes

Induced pluripotent stem cell (iPSC)-derived human astrocytes prepared from a middle age healthy donor (iCell Astrocytes 2.0) and ad-hoc serum-free chemically defined medium (CDM) were both provided by Fujifilm Cellular Dynamics (Madison, WI). Astrocytes were plated on 8-well Millipore Millicell EZ slide chambers (Sigma-Aldrich) coated with 0.06 mg/ml growth factor-reduced basement membrane Matrigel (Corning, Glendale, AZ), and maintained for 1 week in CDM, at 37°C under 5% CO_2_. For evaluation of pro-inflammatory cytokine effects, astrocytes were then cultured for 24 hours in CDM alone (controls) or CDM supplemented with TNF-α (30 ng/ml), IL-1β (10 ng/ml), or IL-6 (20 ng/ml). At the end of the incubation, the media were collected and used as indicated below under “dot blot analysis” to determine the concentration of secreted Nociceptin. Correlation with astrocyte morphology and reactivity in response to cytokine treatments was evaluated by immunostaining with anti-GFAP antibody and confocal microscopy as indicated below.

### Experimental autoimmune encephalomyelitis (EAE)

EAE induction was carried out as previously reported ^39^ with minor modifications. Mice were housed in a controlled 1:1 light:dark cycle room and acclimated to the experimental handlers and room environment for 2 weeks prior to EAE induction. Two different groups were utilized to determine the effects of aging on EAE pathology: 3-4-month-old mice (Younger) and 8.5-9-month-old mice (Older). Prior to injection, animals were anesthetized by inhalation of 3% isoflurane using an ad hoc vaporizer. EAE induction was initiated by administering 100 µg of myelin oligodendrocyte glycoprotein 35-55 peptide (MOG _35-55_) emulsified in complete Freund’s adjuvant (CFA) (500 mg M. Tuberculosis in Freund’s adjuvant) to a final 100 µl volume. Fifty µl of this peptide suspension were injected subcutaneously over each shoulder. This was immediately followed by intraperitoneal (IP) injection of 300 ng of Pertussis Toxin (PT) in 200 µl phosphate buffer saline (PBS). Forty-eight hrs. after MOG_35-55_/CFA and PT administration, a second PT injection was given at the same initial dosis. Control animals were similarly treated, except that the CFA suspension lacked MOG_35-55_ peptide. Animals were daily weighed and scored for clinical symptoms for at least 35 days while blinded to the identity of experimental groups. Clinical symptoms were based on a 0-5 scale: (1) partially limp tail, righting reflex loss, (2) fully limp tail, abnormal gait, (3) hind limb weakness, (4) hind limb paralysis, (5) moribund state. Animals reaching a score of 2 were switched to Diamond Dry Cellulose bedding, and those over the score of 3 were given IP injections of 30% dextrose in saline solution and a gel diet (DietGel 76A, ClearH2O, Westbrook, ME) to ensure adequate hydration.

### Treatment with Nociceptin receptor (NOR) antagonist

Downregulation of Nociceptin signaling in mice was carried out by utilizing BAN-ORL-24 (BAN), a highly specific NOR antagonist. At the first documented EAE clinical score above 0, animals were daily administered a 100 µl IP injection of BAN at a concentration of 1mg/kg body weight in saline solution. Non-treated EAE animals were similarly injected daily with saline vehicle solution alone.

### Nociceptin Analysis

Nociceptin concentrations in human astrocyte-derived cell culture media, human CSF samples, and mouse brain tissue were determined by dot blot analysis as previously described ^28; 37^ with minor modifications. For preparation of mouse samples, the brains from controls and EAE animals were rapidly dissected out and homogenized in ice-cold 10mM Na_2_HPO_4_, 2.7mM KCl, 137 mM NaCl, pH 7.4 (PBS) containing a protease and phosphatase inhibitor cocktail. Adequate aliquots of human CSF, cultured human astrocyte-conditioned media, or mouse total brain homogenates were then spotted onto nitrocellulose membranes. Nociceptin standards were included in the same membrane for quantitative analysis of the samples. Membranes were then air dried for 2 hrs., followed by rehydration in PBS, and 1 hr. incubation in blocking solution (3% nonfat-dry milk, 0.05% Tween-20 in PBS). After 1 hr. incubation with anti-Nociceptin antibody (dil. 1:30,000) in blocking solution, the membranes were washed, re-blocked, and incubated for 1 hr. with the appropriate HRP-conjugated antibody (dilution 1:5,000). After two washes in PBS with 0.05% Tween-20 and four 10 min. washes in PBS alone, membranes were incubated with SuperSignal West Dura chemiluminescence reagent. Detection of immunoreactive dots was then carried out using an Azure Bio-Rad Imager. To correct for potential background level differences, each assay was accompanied by a parallel membrane equally seeded with samples but incubated in the presence of secondary antibody alone. Images were then analyzed by using the NIH ImageJ program.

### Fluorescent immunostaining

For brain and spinal cord immunolabeling studies, mice were anesthetized by IP administration of 2.5% tribromoethanol and transcardially perfused with PBS (3 min.) followed by 4% paraformaldehyde-containing PBS (10 min.), as previously described ^35^. Brains and spinal cords were then dissected out and maintained refrigerated for 1 day in 30% sucrose in PBS. The tissue was then frozen at -80°C in OCT and cut with a cryostat into 20-micron thick slices that were then placed onto microscope slides. For immunolabeling of astrocytes with anti-GFAP antibody, antigen retrieval was performed by placing the slides in pre-warmed citrate buffer (10mM sodium citrate, 0.05% Tween-20, pH 6.0) at 37^0^C for 30 min. For myelin staining with anti-MBP antibody, antigen retrieval was carried out by immersion of the slides in acetone at -20^0^C for 10 min. Prior to staining, all slides were rinsed with PBS and incubated for 1 hr. with PBS containing 5% normal goat serum and 0.3% Triton X-100 (blocking solution). Samples were then incubated overnight with the following appropriate primary antibodies diluted at the indicated concentrations in blocking solution: anti-GFAP (1:500), anti-MBP (1:20), anti-NG2 (1:500), anti-NOR (1:50), or anti-PDGF-R alpha (1:250). After rinsing with PBS and re-blocking for 30 min., samples were incubated for 2 hrs. with the appropriate fluorescently labeled secondary antibodies at a 1:500 dilution, except for anti-rat secondary which was used at 1:250 dilution. For double labeling with rabbit anti-NOR and rabbit Cy5-conjugated anti-MYRF polyclonal antibodies, tissue was first incubated with anti-NOR antibody followed by goat anti-rabbit Alexa-488 as described above. After extensive rinsing of the secondary antibody, slides were re-blocked and incubated for 3 hrs. with CY5-conjugated anti-MYRF antibody (1:100). For all antibodies, at the end of the last incubation, slides were extensible rinsed with PBS and mounted with ProLong glass antifade with NucBlue solution. All fluorescent images were acquired using a Zeiss 880 Confocal Microscope. Maximum projection images were generated from the z-stacks, ensuring no fluorescence crossover between channels.

### Western Blot Analysis

Western blot analyses were carried out as previously described with minor modifications ^37^. Total brain homogenates were prepared in ice-cold PBS supplemented with a protease and phosphatase cocktail and adequate aliquots were then solubilized in Laemmli buffer [60 mM Tris– HCl buffer, pH 6.8; 10% glycerol, 2% sodium dodecyl sulfate (SDS), and 5% 2-mercaptoethanol]. Samples were subjected to SDS-PAGE in 12% polyacrylamide Tris-glycine gels and transferred to nitrocellulose (100 V, 30 min.). Non-specific binding was blocked with 3% nonfat-dry milk, 0.05% Tween-20 in PBS (blocking solution) for 1 hr. at room temperature. Membranes were then incubated overnight at 4^0^C with the appropriate primary antibodies in blocking solution at the following dilutions: anti-GFAP (1:5,000), and anti-β-actin (1:5,000). After rinsing, membranes were re-blocked for 30 min. and incubated for 2-3 hrs. in blocking solution with the appropriate HRP-conjugated secondary antibodies. Following extensive rinsing, immunoreactive bands were revealed with SuperSignal West Dura chemiluminescent reagent and detected using an Azure Bio-Rad Imager. Relative density values were determined by using the NIH Image J program and divided by the corresponding β-actin levels to correct for sample loading and potential differences in transfer efficiency.

### Statistical Analysis

Statistical analyses for two group comparisons were performed by nonparametric Mann-Whitney test, unpaired t-test with Welch’s correction. One-way analysis of variance on ranks (Kruskal-Wallis test) was utilized when comparing more than two groups. All analyses were carried out using the GraphPad Prism 10 program (La Jolla, CA). Differences were considered statistically significant when p-values were <0.05.

## RESULTS

### Nociceptin secretion by human astrocytes is stimulated by proinflammatory cytokines increased in MS

We showed before that normal brain development is accompanied by elevated levels of Nociceptin that coincide with high synthesis of this peptide by early both rodent and human astrocytes ^28^. Nevertheless, Nociceptin levels decrease to background values thereafter coinciding with the beginning of the rapid postnatal period of brain myelination ^37^. However, previous studies showed that at least in cultured neonatal rat astrocytes, Nociceptin production can be stimulated by inflammatory mediators ^40^. Here we show that Nociceptin secretion by human astrocytes is elevated by exposure of the cultured cells to IL-1β, IL-6, and TNF-α **(Fig.1 A)**, proinflammatory microglial cytokines that are increased in MS and play crucial roles in the pathogenesis and progression of this disease ^41–43^. As shown in **Fig. 1B**, GFAP staining indicated that while control cells are characterized by small bodies as slender processes, astrocytes in high Nociceptin-secreting cultures exhibited large cytoplasm, complex cytoskeleton and thicker processes consistent with a reactive phenotype.

**Figure 1.**
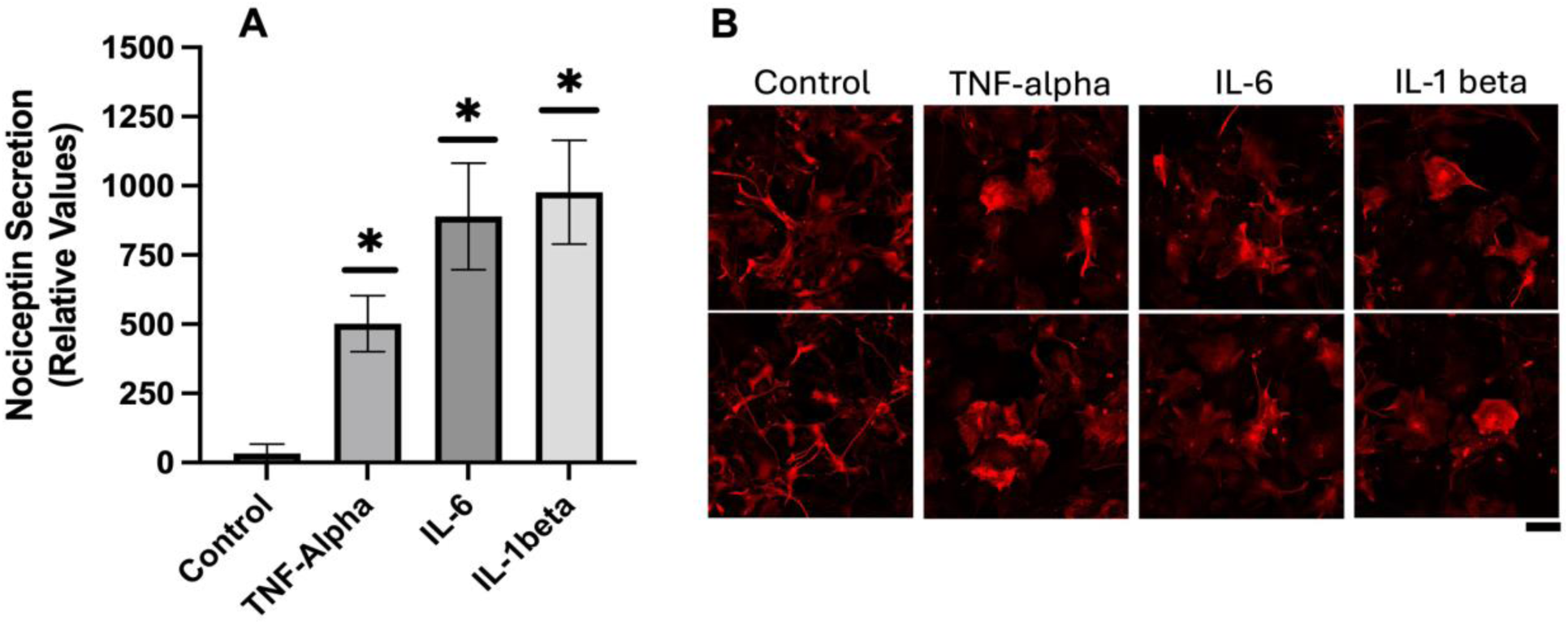
Nociceptin secretion by human astrocytes is stimulated by proinflammatory cytokines increased in MS. **(A)** iPSC-derived human astrocytes were incubated in medium alone (controls), or medium supplemented with TNF-alpha (30 ng/ml), IL-6 (20 ng/ml), or IL-1beta (10 ng/ml). After 24 hrs., Nociceptin secretion was determined in the conditioned media as indicated under Methods. Control vs. TNF-alpha, IL-6, and IL-1beta: * p< 0.05, the bars represent the mean +/- SEM from 3 different cultures/condition. **(B)** Cells were stained with anti-GFAP antibody, shown are 2 representative fields per condition. Scale bar: 50 μm.

### Nociceptin levels are significantly elevated in the cerebrospinal fluid of patients with progressive forms of MS

The preceding observations in cultured human astrocytes led us to question the likelihood of proinflammatory cytokines also stimulating astrocytic secretion of Nociceptin in MS. This possibility was next investigated by first analyzing the levels of this peptide in the CSF from healthy age-matched control individuals and patients with different forms of the disease. As shown in **Fig. 2 A**, low levels and no significant differences were found between samples from age-matched controls and those from RRMS patients. However, the CSF values from patients affected by progressive MS forms indicated Nociceptin levels that are significantly higher than for both controls and RRMS groups.

**Figure 2.**
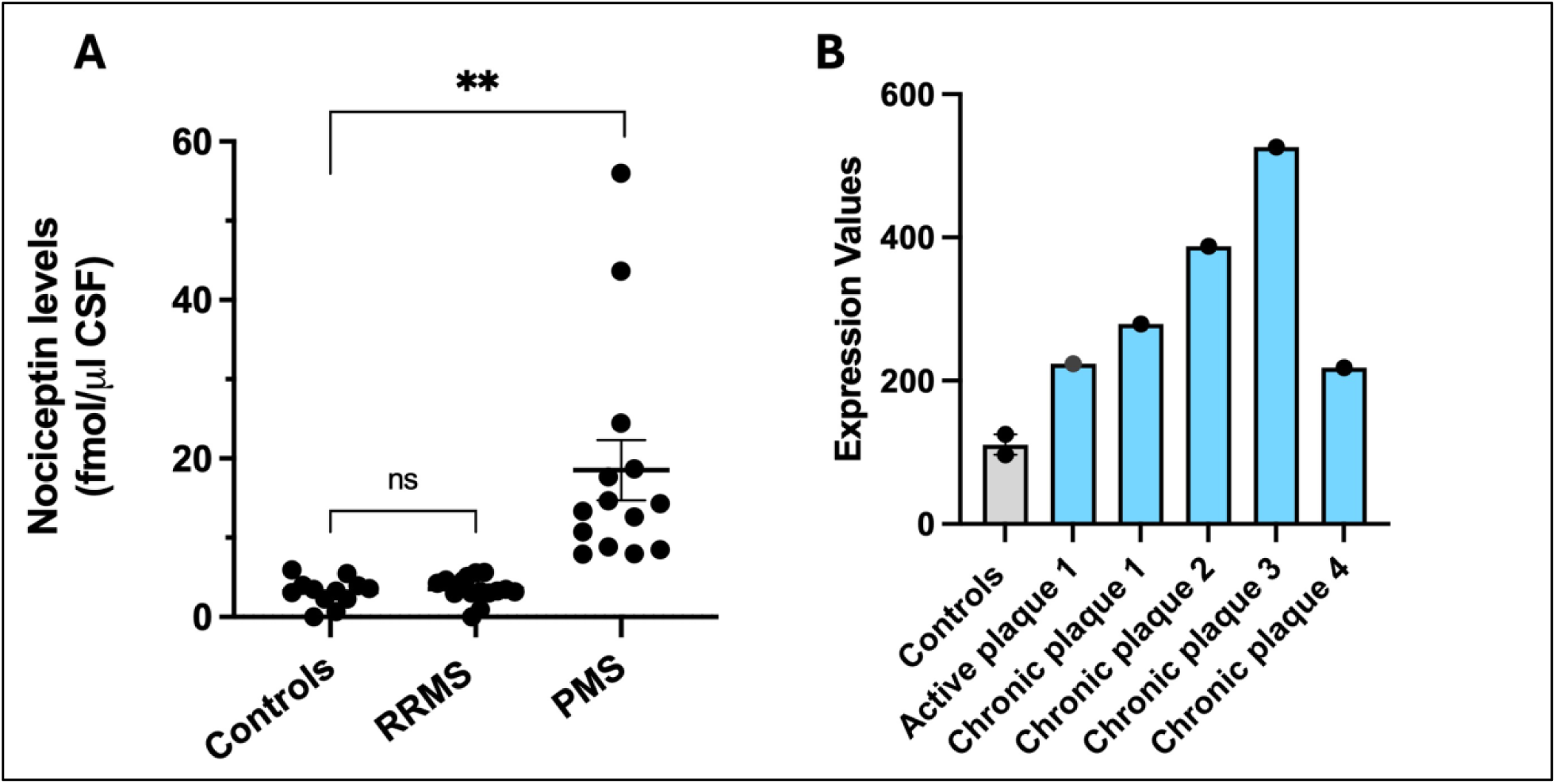
Nociceptin levels are elevated in the cerebrospinal fluid (CSF) of patients with progressive forms of MS, and prepronociceptin (PNOC) mRNA expression is increased in chronic MS plaques. **(A)** Nociceptin levels were determined as indicated under Methods, in CSF from age-matched controls (46.75 +/- 1.59 yrs old, n:12), RRMS patients (45.53 +/- 2.4 yrs old, n:15), and PMS patients (51.07 +/- 1.77 yrs old, n:14). Significance bars indicate SEM. Controls vs. RRMS: ns, p>0.05; PMS vs, Controls and RRMS: ** p < 0.0001. **(B)** The GDS4218 dataset (Han MH et at., 2012) ^44^ from NCBI GEO microarray database was used to analyze PNOC gene expression in controls and MS tissue. Controls (GSM931812 and GSM931813); Active plaque 1 (GSM931814) from MS brain lesion in early stage-active inflammation; Chronic plaques 1 and 2 (GSM931815, GMS931816), chronic active plaques; Chronic plaques 3 and 4 (GMS931817, GMS931818), chronic plaques late stage.

These findings pointed us to explore the NCBI Gene Expression Omnibus (GEO) open-access database that includes a gene expression dataset (GSE38010) from a microarray analysis that identified mRNA from 6,601 targets from two human brain white matter control samples and five MS plaque samples ^44^. Because Nociceptin is generated by proteolytic cleavage of its larger precursor prepronociceptin ^45^, we searched the dataset for mRNA levels transcribed from its encoding gene PNOC. As shown in **Fig. 2B**, PNOC mRNA expression in the five MS samples range from values that are about 2-5 times higher than in the control samples, with the highest levels found in 3 of the 4 chronic plaques. Altogether, these observations raised the question of whether Nociceptin may play a potential role in the remyelination failure that characterizes MS progression ^25^. It is possible to hypothesize that this situation may occur if elevated levels of Nociceptin in PMS recapitulate the inhibitory effect on OLG differentiation that we previously observed during early CNS maturation ^36; 37^.

### Understanding Nociceptin effects using an animal model of MS that takes into consideration the influence of aging in disease progression

Investigating the potential implications of Nociceptin in PMS forms presented the challenge of utilizing an adequate animal model of demyelination/remyelination. One of the most used animal models of MS is experimental autoimmune encephalomyelitis (EAE), in which demyelination is induced by administration of myelin oligodendrocyte glycoprotein 35-55 peptide (MOG_35-55_) to 2-3-month-old mice. This treatment causes a response that results in chronic illness with alternate periods of disease remission, providing an appropriate model for the study of RRMS ^46^. However, this approach was not suitable for the present studies indicating that Nociceptin levels are specifically increased in the progressive forms of MS. Thus, we next decided to use a disease model that takes into consideration the influence of aging in MS progression. Increased age and recurrent episodes of neuroinflammation were previously observed in EAE in Biozzi ABH mice ^47^; and more recently, an interaction between aging with neuroinflammation and demyelination has also been shown in an EAE model induced by adoptive transfer of encephalitogenic CD4^+^ Th17 cells resulting in increased severity and non-remitting disease course in middle-age versus younger C57BL6J mice ^48^. For the present studies, EAE was induced by administration of MOG_35-55_ peptide to 8.5-9-month-old mice, a timing that is consistent with the equivalent human age at which RRMS cases frequently evolve into PMS. As shown in **Fig.3A**, comparison with EAE in younger male and female animals, demonstrates that EAE in older age mice displays significantly higher and persistent clinical scores indicative of severe disease that is more consistent with the situation observed in PMS forms. Importantly, this severe disease course in older mice can be attributed to the response to MOG_35-55_ peptide administration because no clinical symptoms were detected in response to complete Freund’s adjuvant and PT alone **(Supplemental Fig.1)**.

**Figure 3.**
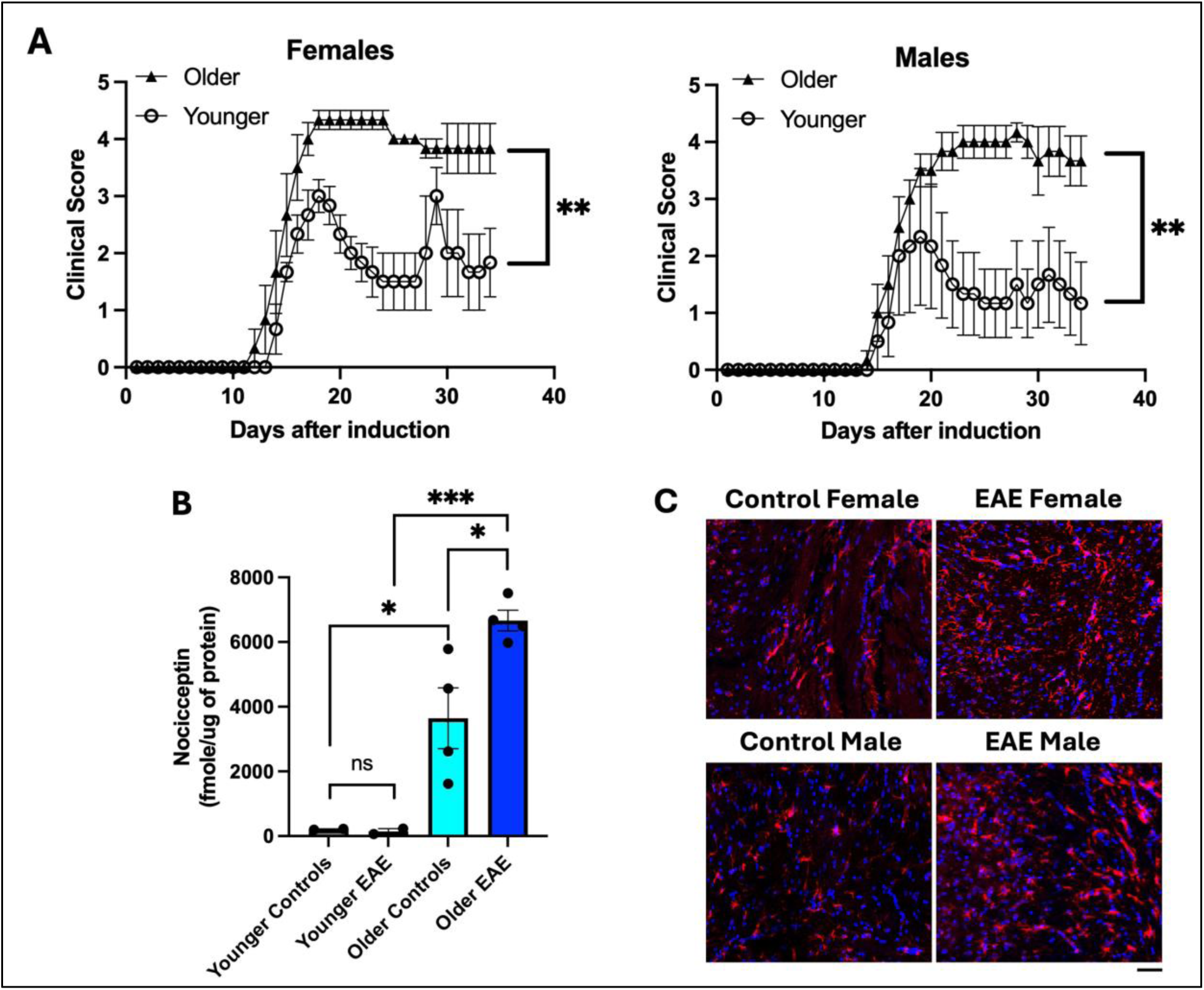
EAE induction in older animals results in persistent high clinical scores and elevated levels of Nociceptin. **(A)** EAE was induced in 8-9-month-old (Older, black triangles) and 3-month-old (Younger, open circles) male and female mice. Clinical Scores: 1-partially limp tail, righting reflex loss, 2-fully limp tail, abnormal gait, 3-hind limb weakness, 4-hind limb paralysis, 5-moribund state. At each time point after induction, the results are the mean +/- SEM from 3 animals/sex. Older vs. Younger, **p<0.001. **(B)** Nociceptin levels in total brain homogenates were determined as indicated under Methods. The bar graph shows the mean +/- SEM, (combined results from 4 animals, 2 males + 2 females, for each age). Younger Controls vs. Younger EAE: ns; Older Controls vs. Older EAE: *p<0.05; Older Controls vs. Younger Controls: *p<0.05; and Older EAE vs. Younger EAE: ***p<0.0001. **(C)** GFAP staining of astrocytes in representative areas in the splenium of the corpus callosum in female and male older controls and EAE mice. Red, GFAP; blue, Hoechst nucleus staining. Scale bar: 50 μm.

Interestingly, comparison between both age groups **(Fig. 3B)** also demonstrated that aging itself is accompanied by increased Nociceptin brain levels, with values in older mice being 3 times higher than in their younger counterparts. Moreover, while EAE at younger age does not increase Nociceptin brain concentrations, levels of this peptide in older mice EAE samples were almost twice as high than in the respective age-match controls and 6 times higher than in both 4-month-old controls and young EAE mice. Consistent with these findings, examination of the corpus callosum following immunolabeling with anti-GFAP antibody also indicated that EAE in older animals results in significant brain increase in reactive astrocytes **(Fig. 3C)**.

### Inhibition of Nociceptin signaling in older mice EAE ameliorates clinical symptoms as well as brain and spinal cord demyelination

The potential contribution of Nociceptin to the progression of disease in older mice EAE was next investigated by taking advantage of the use of BAN-ORL24 (BAN), a potent blood brain barrier permeable and highly specific antagonist of the Nociceptin receptor (NOR) ^49; 50^. We showed before that BAN-administration accelerates OLG differentiation and myelin synthesis in the developing rodent brain, supporting the inhibitory role that Nociceptin plays preventing untimely precocious myelination ^37^. In the present studies, older EAE mice were daily administered BAN from the time of disease onset. As shown in **Fig. 4A**, both male and female EAE animals treated with BAN displayed a significant reduction in the severity of clinical symptoms. Female mice in particular, appeared to be more responsive to the treatment, including regaining their ability to move the hind limbs. On the other hand, EAE controls treated with saline solution vehicle alone exhibited a large increase in clinical symptoms that persisted throughout the studied period. Furthermore, assessment of myelination by staining with antibody against myelin basic protein (MBP) **(Fig. 4B)** and comparison with controls **(a and b)**, indicated that EAE induction in older female and male mice **(c and d)** results in significant demyelination of the corpus callosum that is nevertheless ameliorated by BAN treatment **(e and f)**. Importantly, similar positive BAN treatment effects were also observed when myelination was analyzed in the spinal cord **(Supplemental Fig. 2)**. In contrast with the results in older EAE animals, treatment with BAN does not affect the course of EAE when induced in 3-month-old mice (Supplemental Fig. 3); a finding that is consistent with the negligible levels of Nociceptin at this earlier age (Fig.3B)

**Figure 4.**
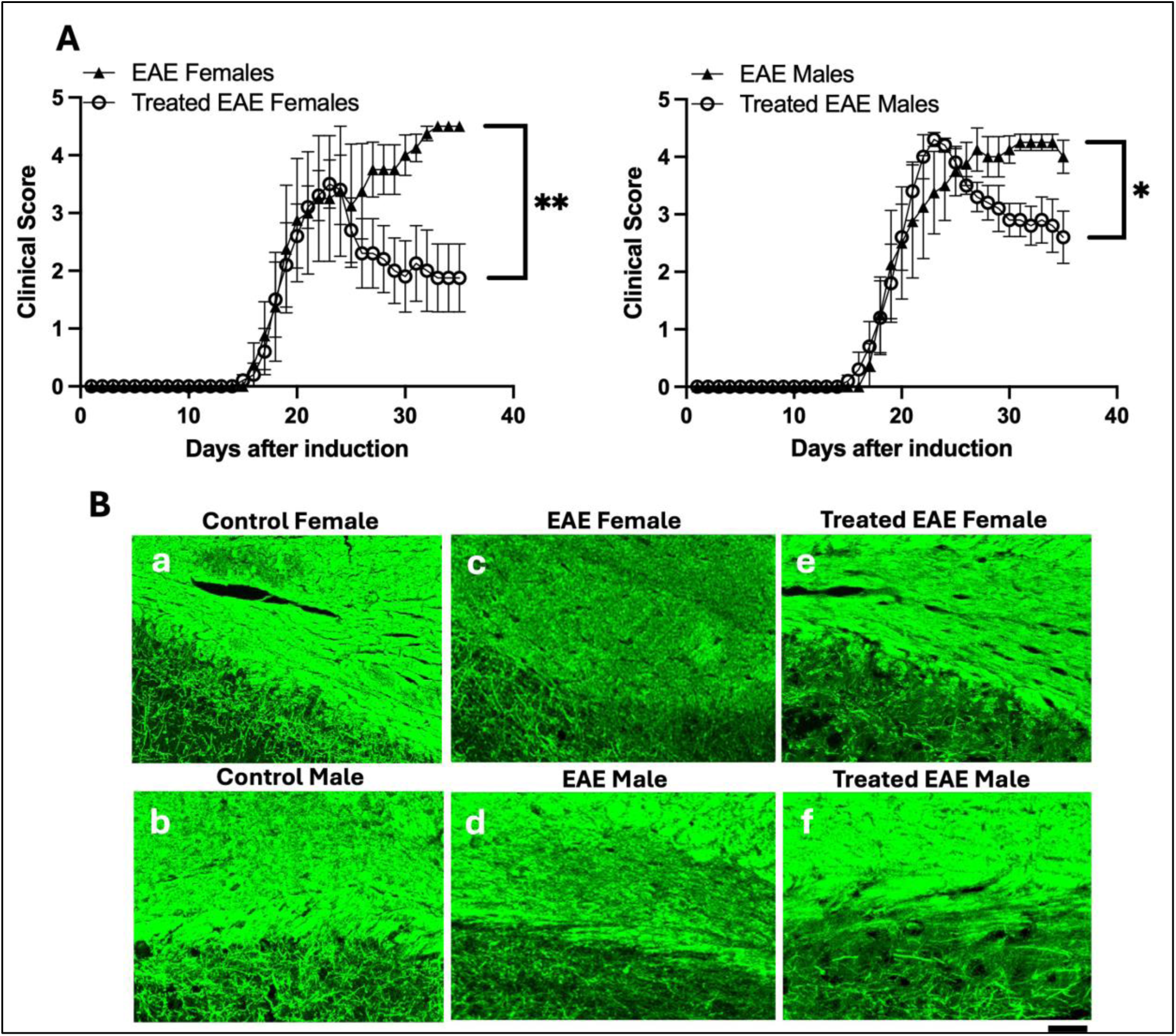
Treatment with the Nociceptin receptor antagonist BAN-ORL-24 ameliorates EAE clinical symptoms and brain demyelination. EAE in older female and male mice was induced by MOG_35-55_/CFA and PT administration as indicated under Methods. **(A)** EAE treated mice (open circles) were daily injected with BAN-ORL24 (1 mg/kg/day, IP) from onset of symptoms as described under Methods. EAE (black triangles) were equally injected from onset of symptoms with saline vehicle alone. At each time point, the results are the mean +/- SEM from 5 animals/sex/condition. Female EAE vs. Treated female EAE: **p<0.001. Male EAE vs. Treated male EAE: *p <0.03. **(B)** The panels show myelination in the splenium of the corpus callosum as assessed by staining with anti-MBP antibody of brain tissue from: **(a and b)** Control mice injected with CFA suspension and PT alone, **(c and d)** EAE, and **(e and f)** BAN-ORL24 treated EAE. Scale Bar 50 μm.

### Decreased clinical symptoms and enhanced myelination in BAN treated older mice EAE is accompanied by changes in the pools of myelinating OLGs

The positive effects of BAN treatment described above pointed to the possibility of elevated levels of Nociceptin in older mice EAE exerting the same inhibitory effects on OLG maturation observed in the developing brain ^37; 38^, a situation that would obstruct remyelination in the adult CNS. To examine this problem, we next investigated the expression of the Nociceptin receptor NOR along the oligodendroglial lineage in the older mice. We found before NOR expression in pre-OLGs ^37^, and here analysis of the corpus callosum of 9-month-old controls and EAE mice **(Fig. 5)** revealed the presence of NOR in OLG progenitor cells (OPCs) identified by their expression of NG2 proteoglycan ^51^ **(Fig. 5A)** and PDGFR alpha ^52^ **(Fig. 5B)**. This finding is particularly important because remyelination is attributed to proliferation and differentiation of OPCs ^53^, but their proliferative and differentiating activity is thought to decrease with aging ^54^. Thus, the presence of NOR in OPCs in the aging CNS raises the question of whether effects of Nociceptin on these cells could play a critical role in the lack of remyelination observed in PMS forms. Supporting this possibility, close examination of OPCs in the corpus callosum of older EAE mice **(Fig. 5 C and D)**, indicated the presence of cells that are mostly rounded and devoid of processes. However, this is in remarkable contrast with the highly branched OPCs present in the EAE animals treated with BAN, a morphology that is reminiscent of that observed in the younger developing CNS where differentiation of these cells leads to myelination. Importantly, the transition of pre-OLGs into differentiating and myelinating OLGs it is marked by their expression of the transcription factor MYRF ^55; 56^ and as shown in **Fig. 6A**, these cells also express NOR. Similarly, NOR expression in OPCs and differentiating/myelinating OLGs is also observed in the spinal cord **(Supplemental Fig 4 A and B)**.

**Figure 5.**
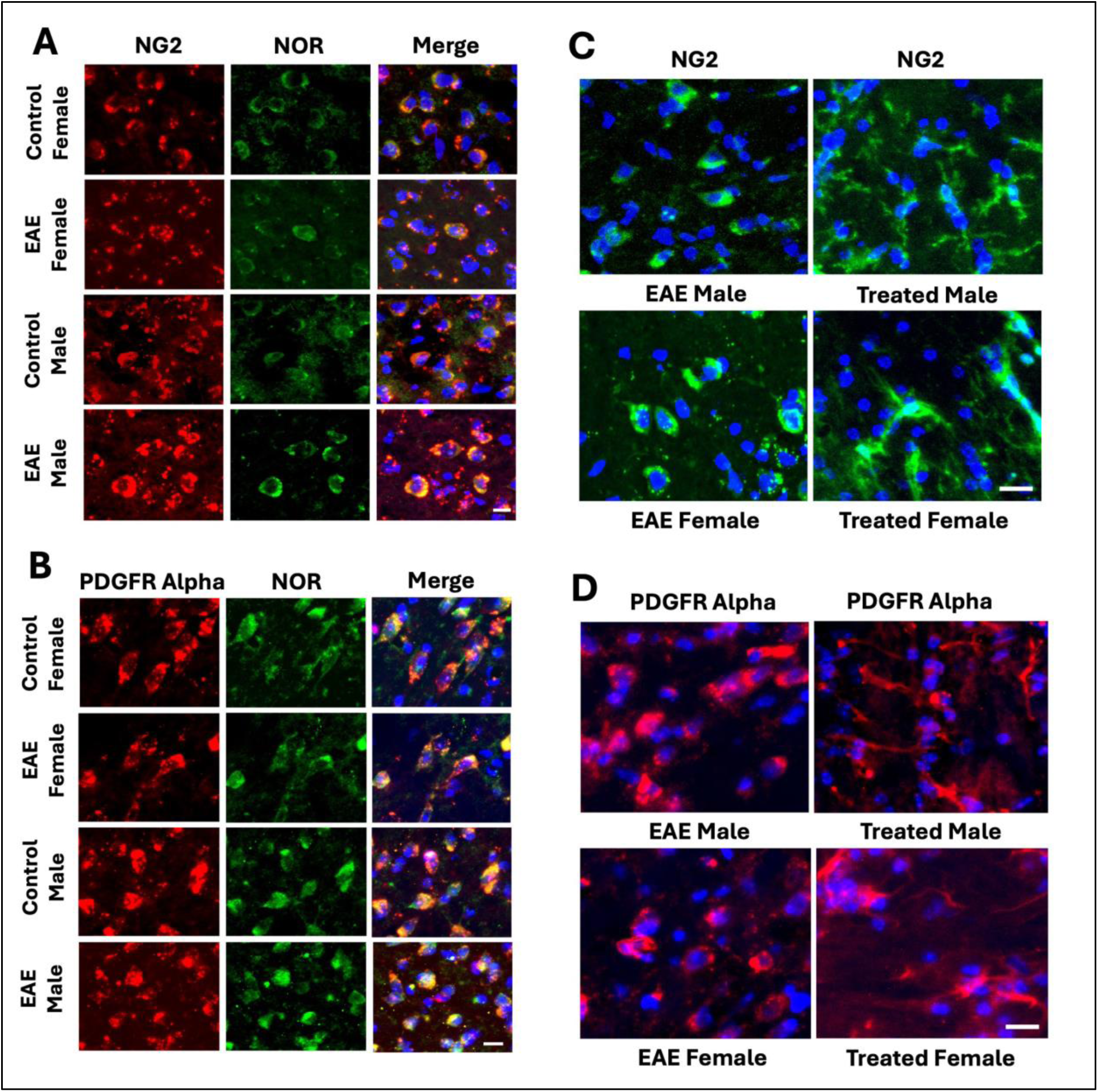
Oligodendrocyte progenitors (OPCs) in older animals express NOR and show increased branched morphology after BAN-ORL 24 (BAN) treatment. Brain samples from controls and EAE female and male mice were subjected to double immunofluorescent staining with anti-NOR antibody to assess NOR presence in OPCs identified by their expression of **(A)** NG2 proteoglycan and **(B)** PDGFR alpha. Shown here are OPCs in comparable areas of the corpus callosum. Scale bar: 10 μm. Staining with both **(C)** NG2 and **(D)** PDGFR alpha shows that EAE brain samples exhibit OPCs with rounded cell bodies devoid of processes. This is in contrast with the branched morphology of these cells in BAN-treated animals. Scale Bar 10 μm.

**Figure 6.**
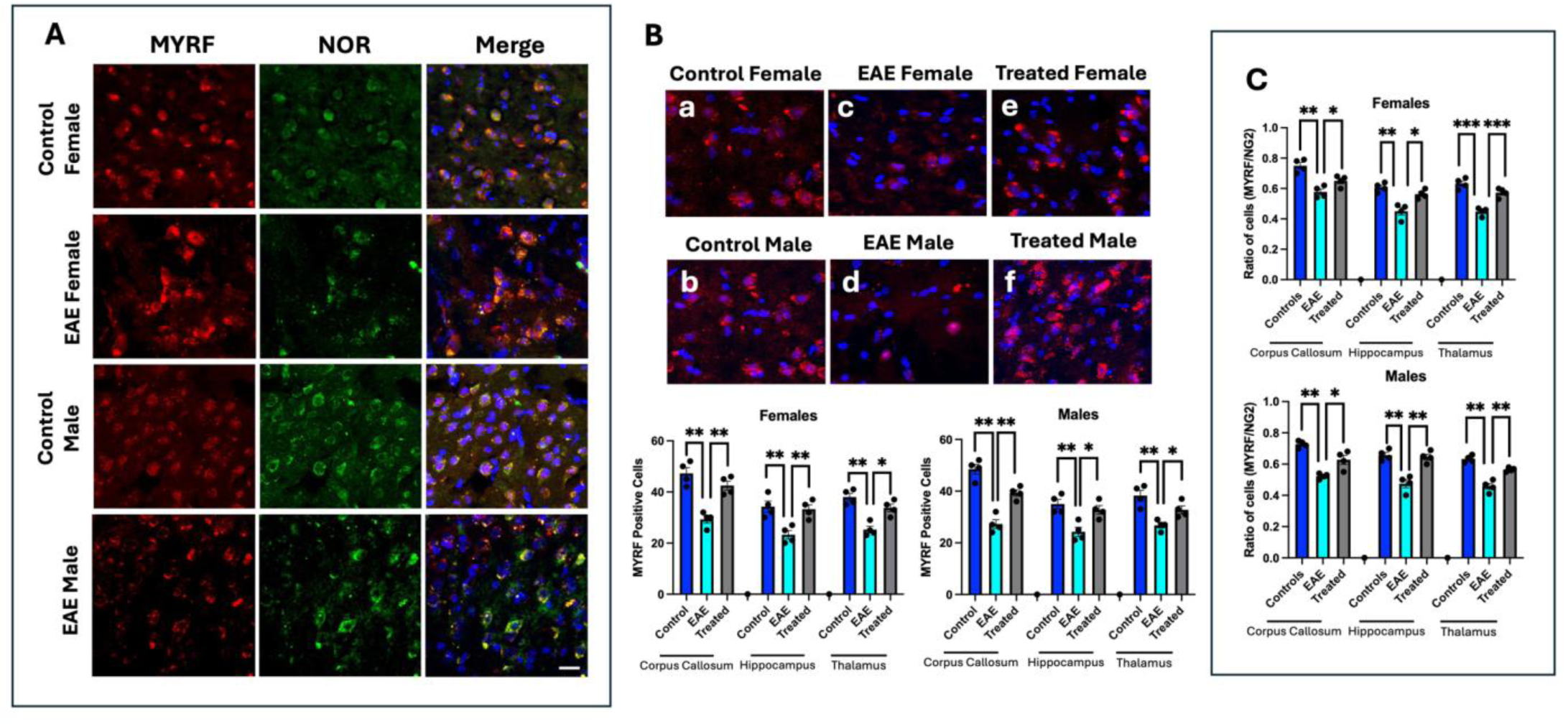
Differentiating/myelinating OLGs in older animals express NOR, and their reduced numbers in EAE are increased following treatment with BAN. **(A)** Brain tissue from controls and EAE female and male mice were subjected to double immunofluorescent staining with anti-NOR antibody to assess NOR presence in differentiating/myelinating OLGs identified by their expression of the transcription factor MYRF. Shown are representative images corresponding to the splenium region of the corpus callosum. Scale bar: 10 μm. **(B)** The panels show MYRF^+^ in control **(a and b)**, EAE **(c and d)** and EAE treated animals **(e and f)**. The bar graphs indicate the number of MYRF^+^ cells in comparable areas of the corpus callosum, hippocampus, and thalamus. The results for Controls, EAE, and EAE treated mice are the number of cells per field expressed as the mean +/- SEM from at least 4 fields/animal from 4 animals/sex/condition. **(C)** The bar graphs show the ratios of MYRF^+^/NG2^+^ cells expressed as mean +/- SEM from at least 4 fields per animal, from 4 animals/sex/condition corresponding to the same areas as indicated in **(B),** Scale Bar 10 μm. *p<0.05, **p<0.01, ****p<0.001).

Quantification in different brains areas (corpus callosum, hippocampus, and thalamus) showed that EAE in older mice results in a significant decreased in the number of MYRF^+^OLGs that is however restored to control levels upon treatment with BAN **(Fig. 6B).** Furthermore, analysis of ratios of MYRF^+^OLGs / NG2^+^OPCs **(Fig. 6C)** revealed that BAN administration results in a significant increase in the proportion of MYRF^+^OLGs, an observation that supports the notion that elevated Nociceptin signaling in aging may inhibit the differentiation of OPCs into OLGs capable of remyelination.

### Reactive astrocytes in the adult CNS re-express NOR

We showed before that early brain rodent and fetal human astrocytes not only produce Nociceptin but also express NOR. However, analysis of rodent brain maturation indicated that the presence of astrocytic NOR is developmentally regulated and restricted to the first postnatal week becoming undetectable thereafter ^28^. Unexpectedly, **Fig. 7A** shows that NOR is re-expressed in GFAP^+^ hypertrophic reactive astrocytes present in both older controls and EAE mice. Furthermore, while these cells are highly increased in both older male and female EAE animals, their numbers are significantly lower upon BAN administration **(Fig. 7 B and C)**. This effect is reflected in parallel significant changes in GFAP levels detected by western blot analysis of total brain homogenates from controls, EAE, and BAN-treated EAE mice **(Fig. 7 D)**. Importantly the re-expression of NOR in GFAP+ astrocytes is also observed in the spinal cord **(Supplemental Fig. 4C).**

**Figure 7.**
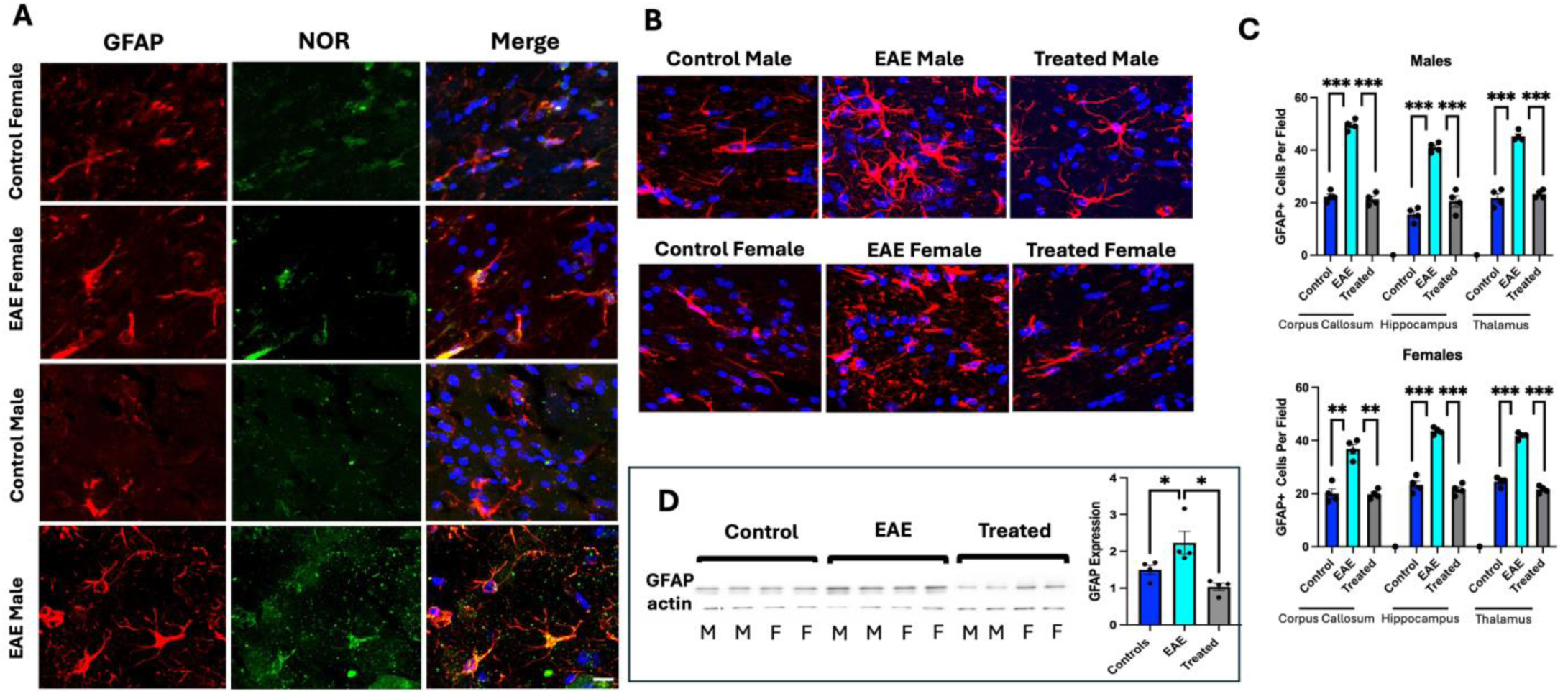
Brains of older EAE mice show re-expression of NOR on GFAP positive astrocytes, and a significant increase in hypertrophic reactive astrocytes which is decreased after BAN treatment. **(A)** Brain samples from controls and EAE female and male mice were subjected to double immunofluorescent staining with anti-NOR antibody to assess NOR presence in astrocytes labeled with anti-GFAP antibody. The panels correspond to representative comparable areas of the splenium region of the corpus callosum. Scale bar 10 μm. **(B and C)** GFAP staining indicates increased numbers of hypertrophic reactive cells in EAE that are reduced by treatment with BAN. The corresponding bar graphs show the number of GFAP positive cells/field expressed as mean +/- SEM from at least 4 fields per animal, from 4 animals/sex/condition. *p<0.05, **p<0.001, ****p<0.0001). **(D)** Western blot analysis of GFAP levels in total brain homogenates from Controls, EAE and Treated animals. The bar graphs represent the mean +/- SEM from 2 males (M) and 2 females (F) per condition. *p<0.05.

Altogether, the findings in CSF and tissue from MS patients and older EAE animals, suggest that Nociceptin may be a crucial player in the age-related progression of MS by not only interfering with OLG maturation and remyelination, but also perhaps further exacerbating neurological dysfunction by affecting different astrocyte populations. The capacity of proinflammatory microglial cytokines to upregulate Nociceptin secretion by human astrocytes, also suggests that this peptide may be an important mediator of the microglial-astrocyte interactions that support MS progression with aging.

## DISCUSSION

The search for efficient strategies to prevent the progression of MS represents a still unmet challenge. Underscoring this problem is the disease multifactorial complexity and the lack of clear understanding of the cellular and molecular mechanisms leading to both demyelination and re-myelination failure. Together with our previous observations, the present findings point to Nociceptin as one of the crucial molecules precluding remyelination in PMS. We showed before that elevated levels of this endogenous peptide during early brain development exert an inhibitory effect that prevents premature OLG maturation and untimely precocious myelination ^37; 38^. However, in contrast with this beneficial early regulatory role, later increases of this peptide as reported here in PMS patients may deter remyelination and/or myelin maintenance in the aging CNS. Support for this possibility is provided by the findings in the older mice EAE model which not only exhibits elevated levels of CNS Nociceptin but also responds to inhibition of Nociceptin signaling with a significant decrease of severe clinical symptoms that is accompanied by increased brain and spinal cord myelination.

Importantly, comparison of Nociceptin levels in young versus older EAE mice suggests that significant increases of this peptide are part of the molecular changes that take place in the aging CNS and contribute to the positive correlation between increased age and MS progression risk. In this regard, studies investigating the factors affecting the rate of conversion of RRMS to PMS determined that, independently of the number of early relapses, the risk of MS progression increases with disease duration and is also significantly accelerated by older age at disease onset ^17–19; 57; 58^. Advanced age is also known to negatively affect the remyelinating capacity of damaged tissue ^59^, with MS progression resulting in increasing size of demyelinating areas colocalizing with activated microglia and reactive astrocytes ^60–63^. Similarly, spinal cord homogenates and infiltrates from older EAE animals exhibit elevated levels of proinflammatory cytokines and widespread microglia characterized by upregulation of genes indicative of a reactive age-associated phenotype ^48^. Within this context, proinflammatory cytokines secreted by microglia in the aging brain, including IL-1β, IL-6, and TNF-α, are known to play crucial roles in the pathogenesis of MS and disease progression ^41–43; 64^. Interestingly, the present results showed that these microglial cytokines also have the capacity of stimulating the production of Nociceptin by human astrocytes, a finding suggesting that this peptide could be an important mediator of microglial-astrocyte interactions supporting MS progression with aging and disease duration.

A crucial finding of these studies is the observation that antagonism of the Nociceptin receptor in older mice EAE not only decreases clinical symptoms and enhances myelination but also alters the pools of OLG populations. The presence of the Nociceptin receptor NOR in both NG2^+^ PDGFR alpha^+^ OPCs as well as in differentiating cells already expressing the transcription factor MYRF strongly suggests that, in addition to the previously reported developmental role in pre-OLGs ^37^, Nociceptin effects in the aging CNS extends to even the earlier steps along the OLG lineage. This is particularly important because early findings analyzing samples from patients with chronic MS already suggested a deficient capacity of the OPCs to replenish mature OLGs damaged upon demyelination ^22^; and at least in animal models, pre-existing mature OLGs do not appear to be responsible for remyelination of demyelinating lesions ^65; 66^. Myelin regeneration at early stages of MS is instead attributed to OPCs that, being also present in human normal and MS tissue ^67^, respond to demyelination by proliferating and migrating to demyelinated lesions where they differentiate into mature remyelinating OLGs ^53^. However, while studies in early MS lesions identified subpopulations of differentiating OPCs, these cells were rarely observed in chronic MS ^25^. This persistent presence of immature OPCs suggested that an inhibition of OLG differentiation is an important cause for remyelination failure in progressive MS and aging. Such a situation may involve intrinsic cell senescence mechanisms as well as age-related cellular environmental changes. OPCs continue to proliferate in the adult CNS ^54^, but advanced age reduces their function and differentiating capacity, a conserved effect observed from rodents ^68–70^, to non-human primates ^71^ and humans ^72^.

The present analysis of NG2^+^ PDGFR alpha^+^ OPCs in older mice EAE indicates the presence of cells with reduced processes, a morphology similar to that described before in aging animals ^73^. This is in remarkable difference with the highly branched OPCs present in EAE treated with the NOR antagonist BAN-ORL24 (BAN), a morphology that is reminiscent of that observed in the younger developing CNS where differentiation of these cells leads to myelination. Furthermore, the increased proportion of MYRF^+^OLGs in the treated older EAE mice supports the possibility that elevated Nociceptin signaling in aging indeed inhibits remyelination by blocking the generation of these differentiated committed OLGs.

Nevertheless, the presence of heterogenous OPC populations in MS ^67; 74^ raises the question of whether a Nociceptin-mediated pathway may also be involved in mechanisms affecting OPC function beyond the generation of myelin making-OLGs. This is because transcriptomic analyses identified the expression of genes involved in antigen processing and presentation in OPCs and OLGs from EAE and MS brain tissue ^75^, and following in vivo fate-tracing studies determined that death of OPCs can occur as a result of their capacity of behaving as antigen-presenting cells to cytotoxic CD8T lymphocytes ^76^. Thus, the possibility of Nociceptin also affecting OPC populations with immunological roles cannot be discarded at this time.

An intriguing finding from the current studies in older mice is the re-expression of NOR in reactive astrocytes of the aging CNS. We showed before that early developing astrocytes not only produce Nociceptin but also only transiently express its receptor NOR. Human fetal brain astrocytes express NOR, and analysis of rodent brain showed that the presence of astrocytic NOR is restricted to the first postnatal week but sharply decreases thereafter being undetectable beyond 10 days of age ^28^. Those *in vivo* and *in vitro* studies also indicated that at early stages of astrocyte maturation, Nociceptin plays a crucial paracrine role specifically stimulating the expression of the glutamate/aspartate transporter GLAST/EAAT1. However, such function of Nociceptin in reactive astrocytes in aging and PMS appears unlikely; or even if present, not sufficient to maintain low levels of glutamate in MS where elevated excitotoxic concentrations of this neurotransmitter have been implicated in the pathogenesis of this disease [for a review on glutamate and MS see ^77^]. Astrocytes comprise a heterogeneous population of cells ^78; 79^ that play a still growing variety of functions including the regulation of synapse formation, function, and elimination; their participation in the induction and functional structure of the blood brain barrier; the homeostatic regulation of pH, and neurotransmitter and ionic concentrations; and the secretion of signaling molecules targeting neurons, microglia, and OLGs ^80–85^. Importantly, transcriptomic studies have shown that aging is accompanied by significant changes in astrocytes from different brain regions, including white matter areas that indicate an increase in reactivity and immune related genes, a problem recently reviewed by Labarta-Bajo and Allen ^86^. Thus, the results indicating a dramatic reduction in reactive astrocytes upon BAN treatment of older EAE mice, raise the possibility of Nociceptin effects also playing a role in aging-associated changes in these cells. Nevertheless, the extent to which these changes in astrocyte populations may contribute to disease in these animals is difficult to assert at this time because astrocyte reactivity has been associated to both neuroprotective and neurotoxic effects ^87; 88^.

In summary, the present observations in human samples and the older age EAE mice point to Nociceptin as a crucial player in the age-related progression of MS. This endogenous peptide may not only interfere with the generation of remyelinating OLGs but could also affect MS progression by exerting direct effects on aging astrocytes and being a downstream mediator of astrocyte-microglial interactions. As a member of the G-protein coupled receptor superfamily, the Nociceptin receptor NOR may offer a unique novel target for pharmacological intervention to prevent the devastating effects of MS progression.

## Acknowledgements

Supported by National Multiple Sclerosis Society grant RG-2206-39621 (C S-B), and NIH-NCI Cancer Center Grant P30 CA016059 (Massey Cancer Center Core Imaging Facility, VCU School of Medicine).

## Supplementary Figures

**Figure 1S:**
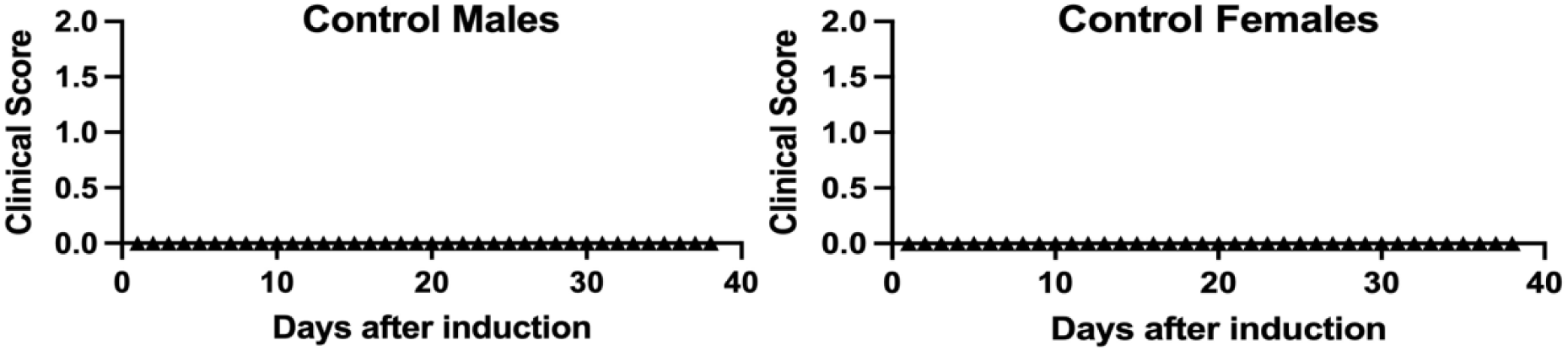
Animals treated with CFA and PT solution alone do not exhibit clinical symptoms of disease. Both control older (8-9 month-old) male and female mice were injected with CFA alone (in the absence MOG_35-55_ peptide) and PT as indicated under Methods. Notice the lack of clinical scores for the entire duration of the study.

**Figure 2S:**
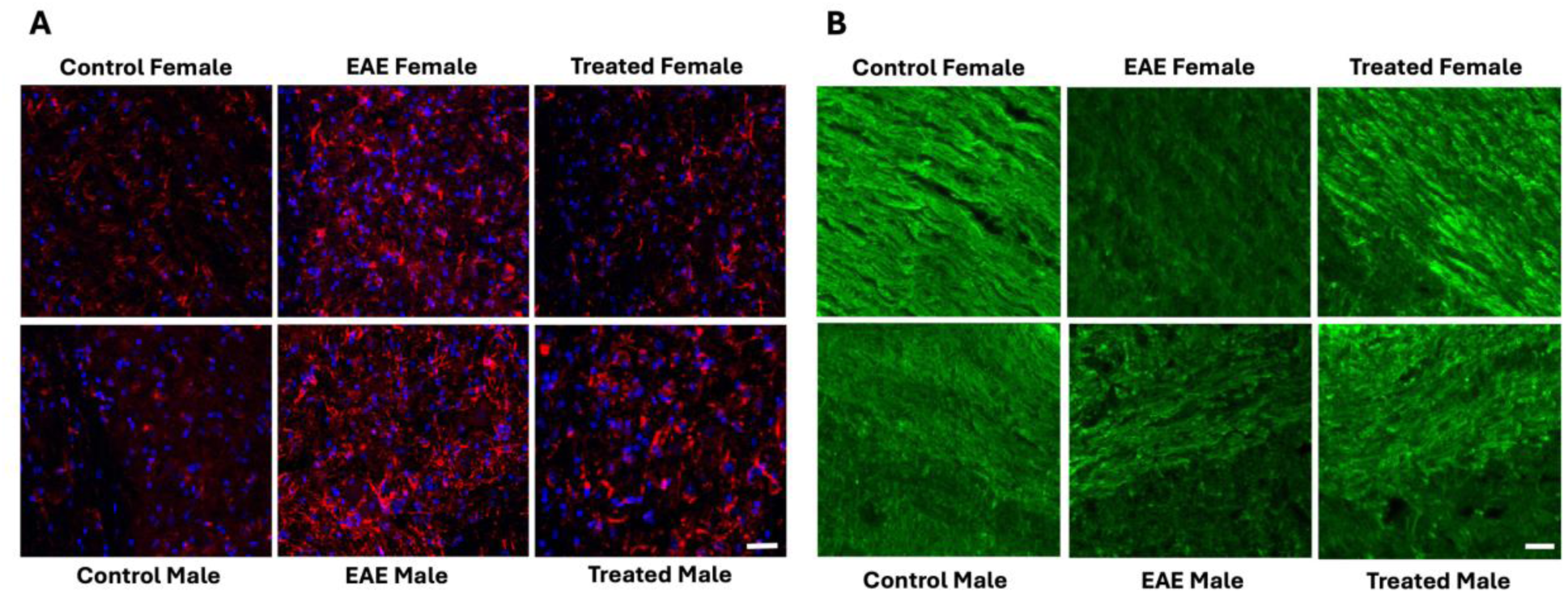
Spinal cords of EAE animals treated with BAN show decreased number of reactive astrocytes, as well as increased myelination. **(A)** GFAP staining of the thoracic spinal cord indicates increased numbers of reactive cells in EAE that are reduced by treatment with BAN-ORL24. **(B)** Thoracic spinal cord sections were analyzed with anti-MBP staining to assess myelination. Scale bar 50 μm.

**Figure 3S:**
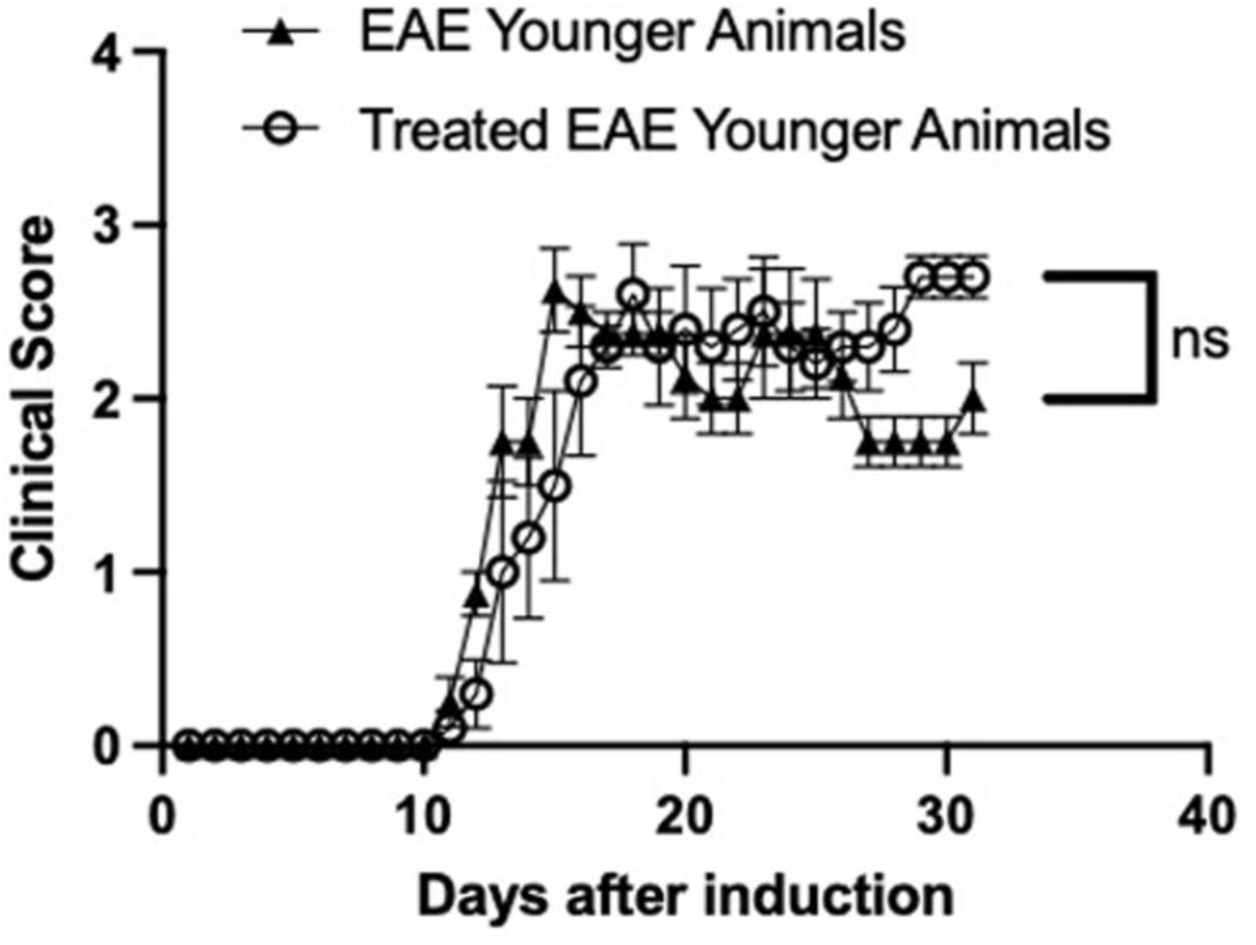
Treatment BAN does not alter EAE symptoms in younger animals. EAE was induced in younger (3-month-old) females by administration of MOG_35-55_ peptide as described under Methods. Mice were daily injected with BAN (1 mg/kg/day, IP) from onset of symptoms as described under Methods. Untreated EAE (open circles) were equally injected from onset of symptoms with saline vehicle alone. At each time point, the results are the mean +/- SEM from 5 animals/condition. No significant clinical scoring differences were detected between treated and untreated EAE mice.

**Figure 4S:**
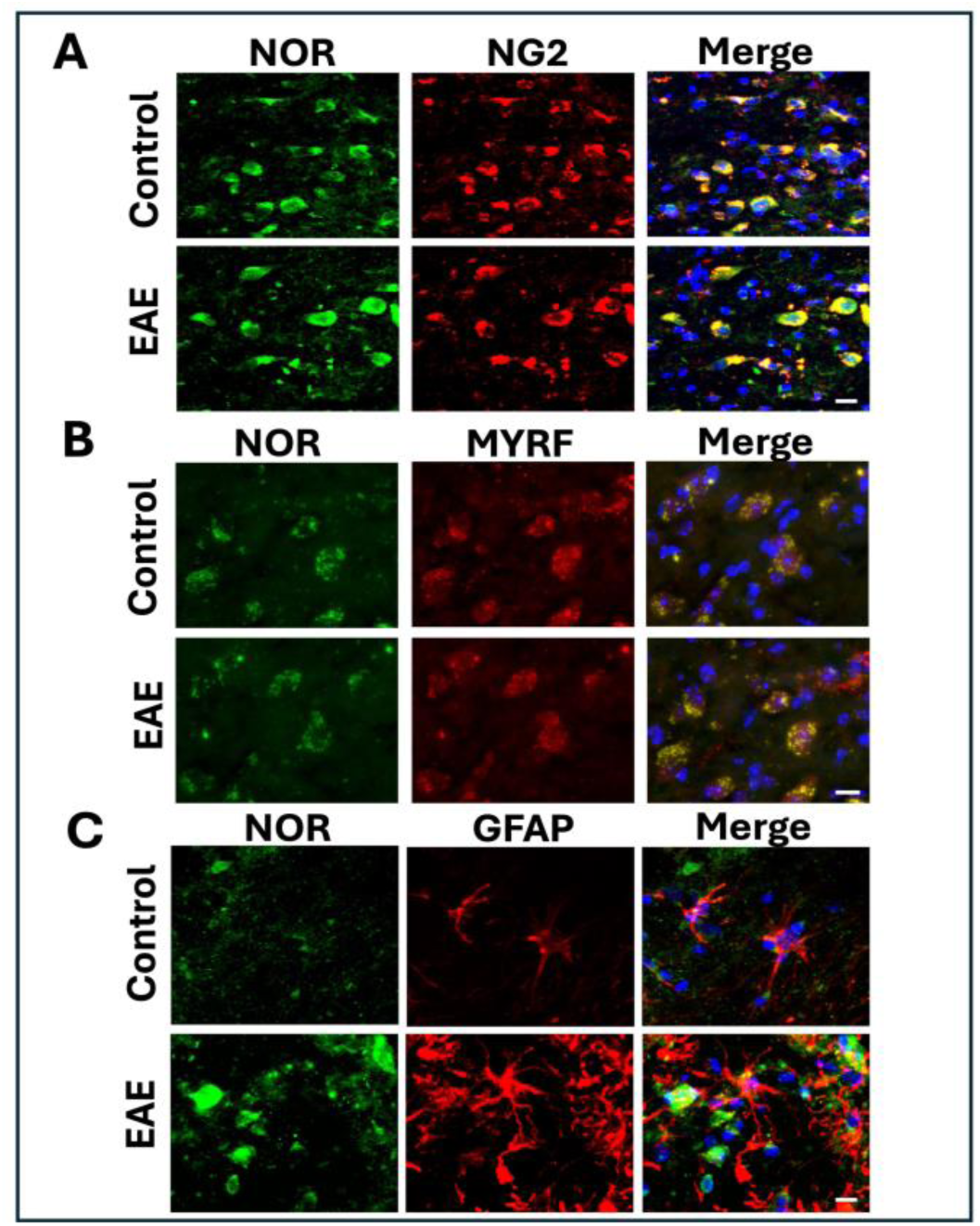
Analysis of NOR expression in astrocytes, OPCs and differentiating/myelinating OLGs. Thoracic spinal cord samples from controls and EAE female mice were subjected to double immunofluorescent staining with anti-NOR antibody to assess NOR presence in **(A)** NG2^+^ OPCs, **(B)** MYRF^+^ cells and **(C)** GFAP^+^ astrocytes. Scale bar 10 μm.

